# Modeling temporal and spatial effects on plant reproduction using observational data

**DOI:** 10.1101/2023.02.20.529237

**Authors:** Flávio R.O. Rodrigues, Cintia G. Freitas, Cristian Dambros

## Abstract

**Background and Aims:** The persistence of a plant species at local sites depend on species ability to survive and reproduce under local conditions. Estimating environmental influence on reproduction is difficult because climatic and soil conditions have synergistic effects on flower and fruit production, and estimating these synergistic effects require tracking a large number of marked individuals. Here, we develop a statistical method that allows investigating the environmental influence on reproduction using observational data only (no marking required).

**Methods:** We surveyed 30 standardized permanent plots on central Amazonia for six herbaceous families of Zingiberales and Poales. The plots were surveyed for twelve consecutive months. Using the newly developed method, we estimated the effect of climatic (temporal) and edaphic (spatial) covariates on flower and fruit production.

**Key results:** We demonstrate that plant reproduction can be estimated from observational data only when enough temporal and spatial data are available. By using the proposed statistical method, we show that the conversion of flowers into fruits in Amazonian monocots is highest in sandy soils, and the difference in conversion rates between sandy and clayey soils is more pronounced in wetter months.

**Conclusions:** By comparing the production of flowers and fruits with previously published data on species occurrence, our results suggesting that species distribution is limited primarily by species capacity to produce fruits (e.g. limited pollination) and not by the capacity to produce flowers. Due to the association of fruit production to climatic and edaphic variables, our results point to potential changes in species reproduction and distribution under future climatic regimes.

## INTRODUCTION

The stable persistence of a species in a site depends on the species to survive and reproduce under local environmental conditions (Giesel 1976). Changes in climate, such as the increase in seasonality and decrease in rainfall, are likely to affect the distribution of many plant species (Kelly and Goulden 2008). However, species will only be able to track changes in climate if soil conditions are appropriate for growth and reproduction (Figueiredo *et al*. 2017; Zuquim *et al*. 2020). Although many studies have investigated how environmental gradients are associated to the presence of plant species (Costa *et al*. 2005; Castilho *et al*. 2006) and reproduction (Freitas *et al*. 2016), very few have determined how climatic and edaphic conditions interact to affect plant reproduction.

Estimating the effect of the environment on flower and fruit production can be difficult because climatic and soil conditions interact in complex ways (Koster *et al*. 2004; Deng *et al*. 2018). Previous studies show environmental factors, such as rainfall (McLaren and McDonald 2005; Lemoine *et al*. 2017) and soil depth (Ågren *et al*. 2008; Valdez-Hernández *et al*. 2010) can control flower production. Likewise, plants produce more fruits with the increase in rainfall (Opler *et al*. 1980) and water availability (Lee and Bazzaz 1982; Akhalkatsi and Lösch 2005). However, the effect of climate and soil conditions on flower and fruit production is likely to be synergistic, and the combined effect of temporal and spatial variables on plant reproduction is poorly known.

To disentangle the direct and joint effects of climatic and edaphic conditions on plant reproduction it is necessary to investigate flower and fruit production simultaneously along climatic and edaphic gradients, using both temporal and spatial replicates. However, studies of plant reproduction with spatial and temporal replicates are rare (but see Teller *et al*. 2014) and only developed with few species. The limited number of studies is likely due to the difficulties in marking and tracking a large number of individuals at multiple sites, which is usually required for data analysis (Rees *et al*. 2014). Also, the flowers of many plants are delicate and can be damaged by handling. To our knowledge, no statistical model exists allowing the estimation of reproductive rates using only survey data and including the effect of temporal and spatial covariates.

We developed a statistical model to disentangle the effect of the environment on flower and fruit production based on field observations and using unmarked individuals only. The low requirements of the model allowed the collection of large amounts of data to determine the effect of a broad range of climatic and edaphic conditions simultaneously on plant reproductive outcome. We quantified the production of flowers and the conversion of flowers into fruits along rainfall and soil texture gradients to determine how these variables affect plant reproduction in a herbaceous community in Amazonia. Rainfall and soil sand content increased the conversion rate from flower to fruits and the probability of fruit persistence. Areas predicted by the model to have higher rates of flower conversion and fruit persistence also have greater abundance and richness of herbaceous plants, suggesting reproduction is a limiting factor for plant distribution and diversity.

## MATERIALS AND METHODS

### Model construction

The number of fruits present in a plot at a given time depends both on the number of flowers produced at a previous time, as well as on the conversion of flowers into fruits and the number of fruits that persists. Therefore, estimating fruit production is more complicated than estimating flower production. Current statistical methods to estimate transitions of flowers to fruits in plant populations require marking individuals (Rees *et al*. 2014) because fruits last for long periods, and the rate of conversion of flowers into fruits cannot be measured simply as the proportion of fruits and flowers observed in consecutive censuses. However, when multiple sites are surveyed repeatedly over time, marking and tracking a large number of individuals is costly and labor-intensive. In addition, many plant species produce small and delicate flowers that are impossible to mark individually or in which manipulation could induce flower mortality.

We developed a specific model to simultaneously estimate the rate of flower conversion into fruits, the rate of fruit persistence from one time-step to the next (eg. consecutive months), and the effects of temporal and spatial covariates (rainfall and soil clay content in this study) on these rates. The model represents an extension of the logistic regression, and the parameters are estimated by maximizing a likelihood function. The model jointly estimates the likelihood of observing a given number of fruits at time t+1 given the number of flowers and fruits observed at time t, the probability of a flower to be converted into a fruit from t to t+1 (p_h=1_), and the probability of a fruit to persist from t to t+1 (p_h=2_). The likelihood function for the model is described as:

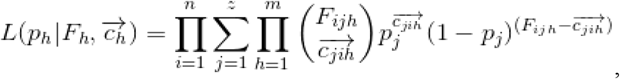

Where i, j, and h represent indexers of site and time (i; e.g. month), the number of fruits or flowers persisting from t to t+1 (j), and the number of sources of fruits (eg. m=2; flowers (h=1) and fruits (h=2)).

To simplify our model, we used only two sources of fruits (m=2): flowers at time t (h = 1), and fruits at time t (h = 2), and the likelihood function can be described by the equation:

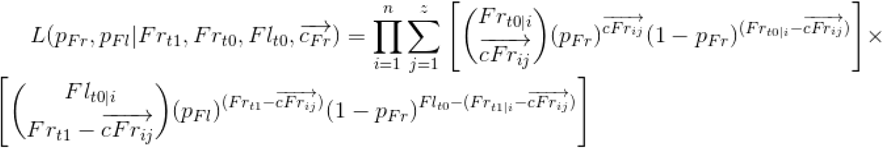

Where pFl is the probability of a flower observed at time t to become an observed fruit at time t+1, pFr is the probability of a fruit observed at time t to persist until t+1. Frt1 is the number of fruits observed in the plot at time t+1, Frt0 is the number of fruits observed in the plot at time t, Flt0 is the number of flowers observed in the plot at time t. cFr is a vector describing all possible combination of fruits at time t that could have been maintained to generate Frt1 fruits at time t+1.

The first term inside the summation describes the probability that an observed fruit persists, whereas the second term describes the probability that a flower is converted into a fruit. The summation sums over the multiple combinations (z) of fruits maintained and flowers converted that could generate the observed number of fruits at t+1. All possible fruits maintained are represented in the vector cFr. Because fruits at time t+1 can only result from fruits present at time t or flowers converted (assuming no detection errors), the possible number of flowers converted is Frt1 – cFr. In our model with two sources of fruits, the vector cFr can be calculated as:

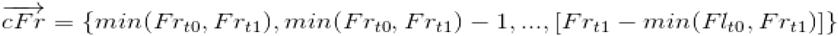

In other scenarios in which the total number of individuals cannot be directly observed (eg. undetected flowers or fruits at time t), cFr could be drawn from a random variable (eg. Poisson or Negative Binomial). In these cases, z would go to infinity and the summation would be weighted by the probability of elements in cFr. However, for simplicity, we assumed all fruits and flowers at time t that could generate fruits at time t+1 were observed. This assumption is reasonable for sessile organisms in a delimited censured area. In the model above, there are only two parameters to be estimated, pFr and pFl. A final step in the model is to include temporal and spatial covariates as predictors of pFr and pFl. As in logistic regression, pFr and pFr were transformed using a logit link function, and the logit-transformed variables represent a linear combination of predictors:

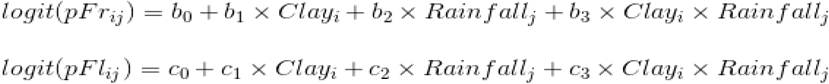

where i and j represent plots i = {1, 2, 3, …, 30} and temporal observations j = {2, 3, …, 9}. Note that if fruits were assumed to not persist from one month to the next, the model would be reduced to the logistic regression (m = 1 in eq. 1). However, this is not the case in most species and is not true for Amazonian herbs because the number of observed fruits at a given month is almost always higher than the number of flowers observed at previous months (Fig. 1). The model presented here could also be easily extended to include other sources of fruits (eg. green fruits, ripe fruits, cross-pollinated flowers, self-pollinated flowers, etc.). All eight parameters in our model (b_0_, b_1_, b_2_, b_3_, c_1_, c_2_, c_3_, and c_4_) were simultaneously estimated by maximizing equation 2 using the Nelder-Mead algorithm (Nelder and Mead 1965).

**Figure 1.**
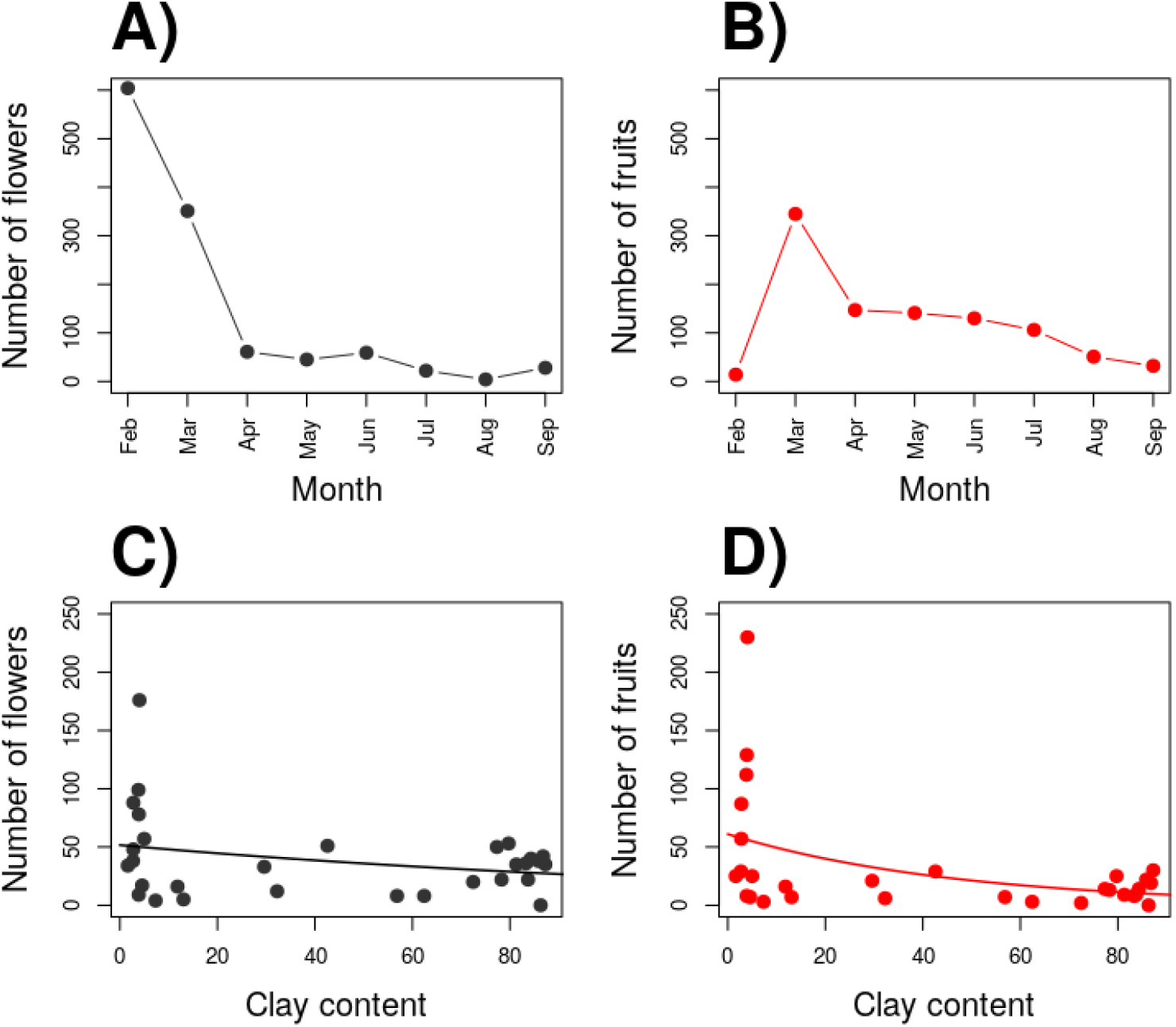
Changes in flower and fruit production over time (A-B) and along the soil clay content gradient (C-D). Except in february and march, all plants had more fruits in a given month than flowers in the previous month (A-B), demonstrating the long-lasting nature of fruits in the investigated species. Flower and fruit production was higher in soils with low clay content (C-D).

To test for the significance of the effect of the covariates on pFr and pFl, subsets of models were created by removing the effects of individual variables (set coefficient to zero; null model). We then refit the model to the remaining parameters and calculated the likelihood for the null model. Finally, we performed a likelihood ratio test (LRT) to compare the full model with individual null models.

In addition to LRT, we simulated the data 999 times under the null model, refit the data generated under the null model, and compared the observed coefficients with those estimated using the simulated data. P-values were calculated as the frequency in which the absolute coefficients in the null model were at least as high as the absolute values of the observed coefficient:

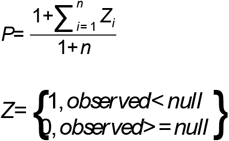

### The case of Amazonian herbs

Data on flower and fruit production of herbs were obtained at Reserva Florestal Adolpho Ducke – RFAD (Ducke forest hereafter) of the National Institute for Amazonian Research. Ducke forest is located 26 km north of Manaus, Amazonas state, Brazil, and covers 10×10 km of terra-firme forest (not subject to periodical flooding) in central Amazonia. The mean annual temperature is 26 °C and air humidity varies from 77 to 88%. The mean annual rainfall is 2362 mm with the driest month in September (Marques Filho *et al*. 1981). The main environmental gradients in the area are associated with topography, soil texture, and water availability. The terrain varies from sandier soils near the drainage in the valleys (less than 10% of clay) to clayey soils in the plateaus (more than 80% of clay; Quesada *et al*. 2011).

### Sampling design and data collection

Ducke forest has a grid of 30 regularly 1-km-spaced plots marked from north to south and east to west, established by the PPBio program (Long term Research Program in Biodiversity; Pezzini *et al*. 2012) and covering a total area of 25 km^2^. Each plot is represented by a 250 m long and 40 m wide line. The central line of each plot follows the elevation contour to minimize variation in altitude and edaphic conditions within the plot.

In each plot, the number of individuals flowering and fruiting of all reproductive monocots (e.g. nineteen herb species and six families of Zingiberales and Poales order) was recorded from February to September 2010. Most species (17 of 19) have small and delicate flowers and flower abortion could be easily increased by manipulation. To increase accuracy and precisely estimate parameters for flower and fruit production under the developed statistical models, we sampled a higher number of plots and individuals per plot than previous studies conducted with plants with larger flowers (e.g. Sakai *et al*. 1999; Kudo and Hirao 2006; Alfaro-Sánchez *et al*. 2017; Pires *et al*. 2018).

Mean monthly rainfall was obtained for the study area from the Research Coordination in Climate and Water Resources (CPCR - INPA). To measure soil clay content, soil samples were collected at six points distant 50 m from each other along the main axis of the plot at a depth of 0–5 cm. The samples were bulked together to produce a composite sample that was cleaned (i.e. roots, twigs or any debris were removed), air-dried and passed through a 2 mm sieve. Clay content was calculated as the percentage of the soil weight after washing off the sand and silt (Manual de Análise Granulométrica do Solo – INPA, 2015 https://ppbio.inpa.gov.br/manuais).

### Data Analysis

To estimate the effect of soil clay content and rainfall on flower production, we used a Generalized Linear Model with a Poisson distribution of errors and included the number of individuals flowering as a response variable, and soil clay, rainfall and the interaction of soil clay and rainfall as predictor variables. We then used the model described in this paper to estimate the rates of flower conversion into fruits and fruit persistence along climatic and edaphic gradients. Soil clay, rainfall and the interaction of soil clay and rainfall were used as predictor variables.

Rainfall and soil clay content are likely to have similar effects for most species included in the study (Costa *et al*. 2005; Rodrigues and Costa 2012). Because many species were too rare for reliable estimates, we determined only the overall rates along the environmental gradients at the community level (i.e. for all species combined). All analyses were performed in the R program (R Core Team 2019).

## RESULTS

We monitored 668 individuals flowering and/or fruiting, representing nineteen species of monocot herbs and six families of the Zingiberales and Poales order. *Heliconia acuminata* was the species with the highest number of individuals flowering (N_fl_ = 11.69, sd_fl_ = 19.53) and fruiting at a given month (N_fr_ = 6.19, sd_fr_= 8.70), along with *Monotagma tomentosum* (N_fl_ = 5.08, sd_fl_ = 11.18, and N_fr_ = 8.46, sd_fr_ = 11.62). The majority of species were observed flowering and fruiting in few plots (n < 10), preventing analyses of flowering and fruiting separately for each species. Despite the low occurrence of other species, the three most common species (*H. accuminata, Ischnosiphon arouma*, and *M. spicatum*) tended to flower and fruit in the same plots (r_flower_ = 0.28-0.42; r_fruit_ = 0.24). Moreover, flowering and fruiting were similar among species (correlation for flowering between species pairs = 0.55-0.7 and correlation for fruiting between species pairs = 0.72-0.79). When all months are combined, the overall number of flowering and fruiting individuals decreased in areas of high soil clay content (b = -0.06; p < 0.03; Fig. 1C;D). Similar results were obtained when using individual data from individual months and including rainfall as a covariate (Fig. 2).

**Figure 2.**
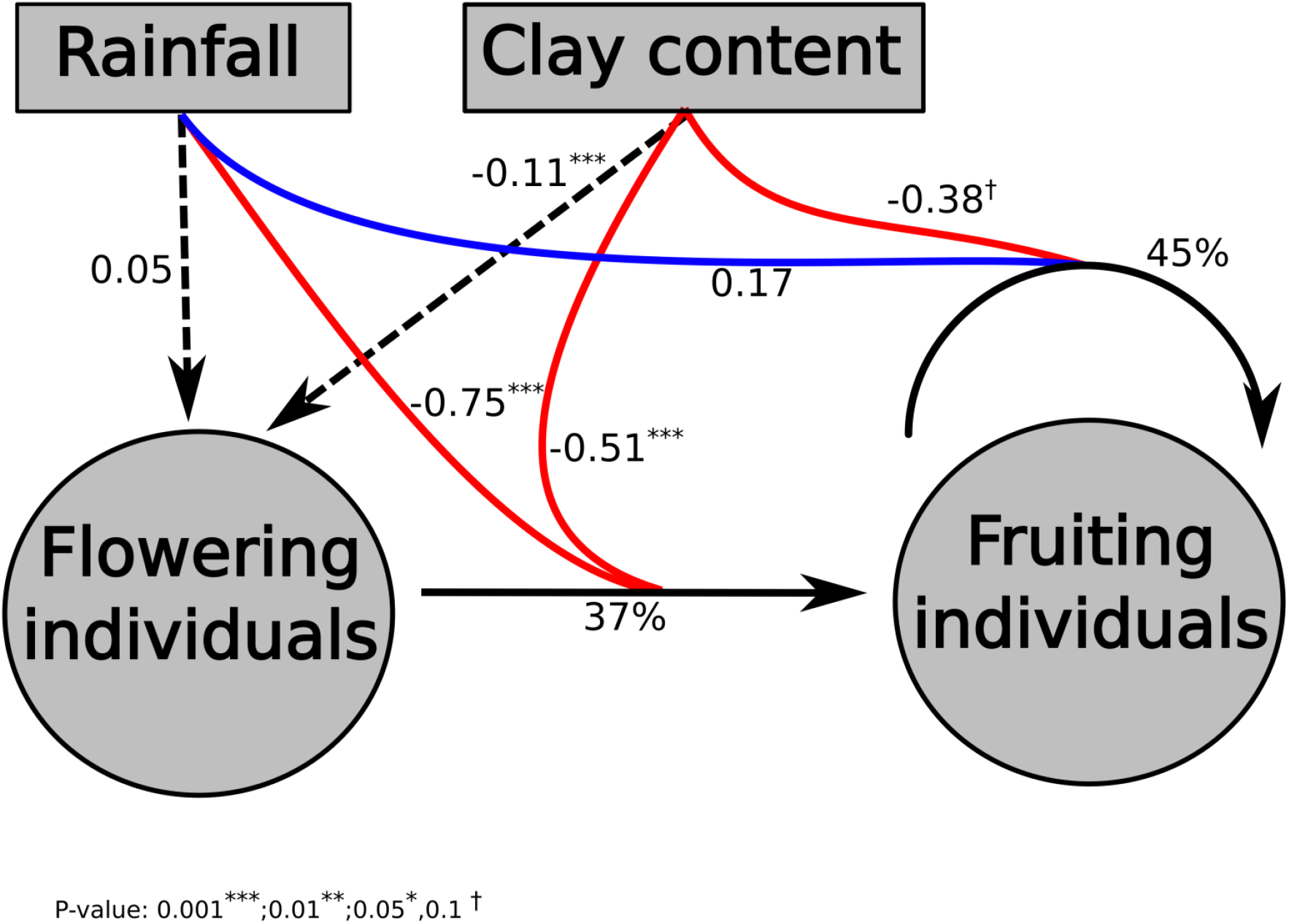
Diagram of all results associating rainfall and soil clay content to flower and fruit production and fruit persistence. Blue and red arrows represent positive and negative associations, respectively. Black arrows are non-significant associations or the intercepts representing the average conversion of flowers to fruits and average fruit persistence (logit transformed).

An average of 37% flowers was estimated to convert into fruits from one month to the next (Fig. 2). However, the rate of flower conversion into fruits was lower in months of higher rainfall (b = -0.75; p < 0.001; Fig. 2; Fig. 3B) and in areas of high soil clay content (b = -0.51; p < 0.001; Fig. 2; Fig. 3B). Around 45% of fruits persisted in a plot from one month to the next (Fig. 2). We have not detected a significant association of fruit persistence with rainfall and soil clay content (Fig. 2; Fig. 3A). Except for February and March (months with highest number of flowers and fruits, respectively), the number of fruits observed at a given month was always higher than the number of flowers at the previous month, indicating that the observed fruits result both from flower conversion and the persistence of fruits from previous months (Fig 1A-B).

**Figure 3.**
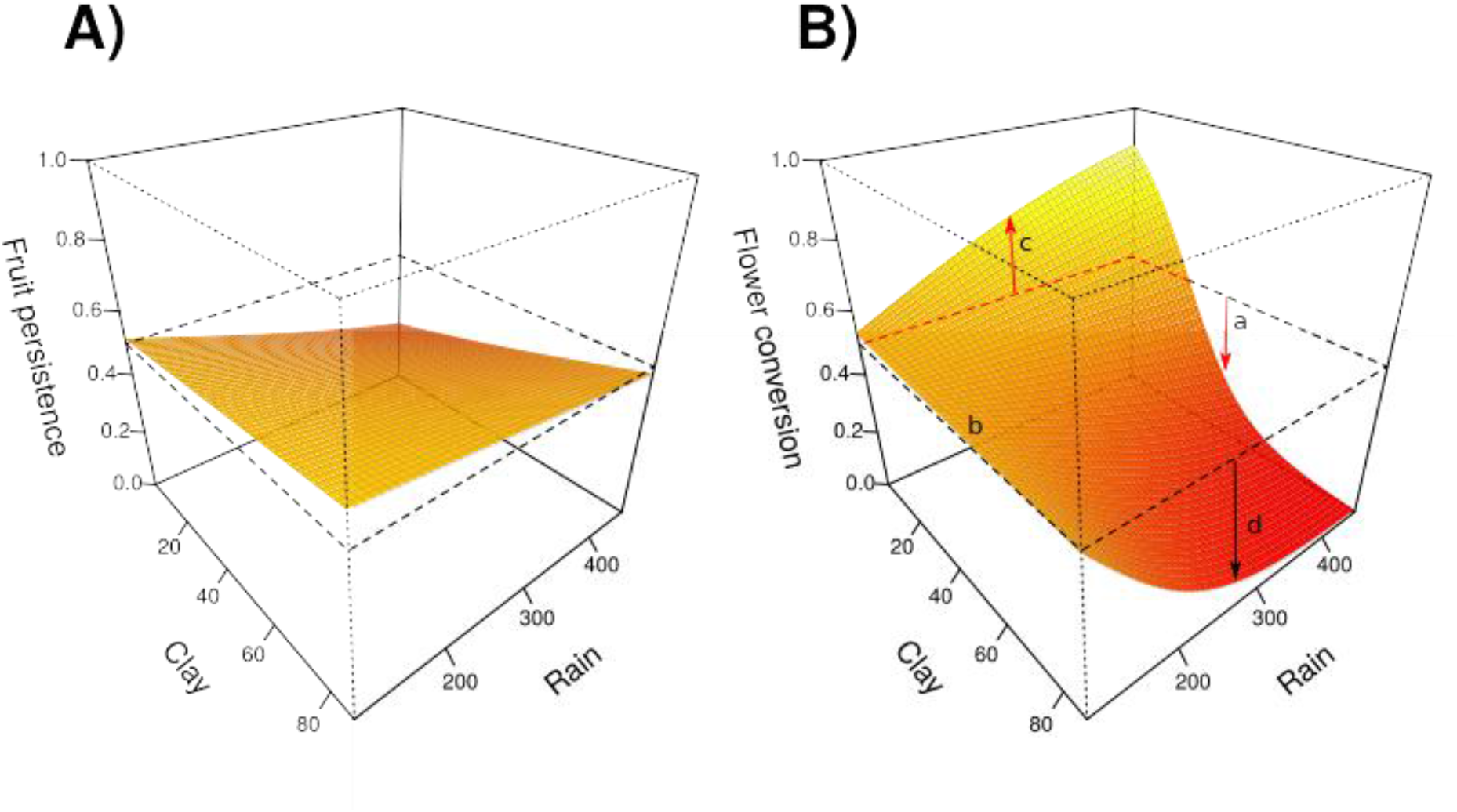
Association of rainfall and soil clay content on fruit persistence rates (A) and flower conversion rates (B). The effect of clay content and rainfall was stronger for flower conversion than fruit persistence. In sandy soils (low clay content), flower conversion increased with rainfall, whereas in clayey soils flower conversion decreased with rainfall.

In addition to the direct effects of rainfall and soil clay content, the effect of rainfall on flower conversion changed from sandy to clayey soils (ie. significant interaction of these variables; p = 0.003). Rainfall had a positive effect on flower conversion on sandy soils and a negative effect on clayey soils (Fig. 3B).

## DISCUSSION

We present two major advances in the evaluation of reproductive success in plants. First, we developed a model able to capture the subtle balance between flower-fruit conversion, fruit persistence, and the effect of the environment on these processes by calculating the probabilities of phase transitions without plant manipulation. This approach is promising when marking individuals is difficult and when the least of human interference is desirable, such as in permanent plots of long-term monitoring programs. Then, by dissociating the effect of climate and soil conditions on fruit production, we demonstrate the effect of climatic variables on plant reproduction cannot be satisfactorily understood without considering the edaphic conditions where reproducing individuals are.

The flowering of monocot herbs in the *terra-firme* forests of central Amazonia was higher in sandy soils and little affected by rainfall. Nevertheless, fewer than half of the flowers became fruits and the rate in which these flowers become fruits is strongly associated with soil clay content and rainfall. Therefore, our results suggest soil sand has a positive relationship with flowering, but soil sand and water availability impose a much stronger restriction on the conversion of these flowers into fruits. Tropical forests are highly productive ecosystems; however, plant growth is known to be limited by nutrients (Santiago *et al*. 2012). Here we extend this evidence to plant reproduction, as seen before for palms (Freitas *et al*. 2016), showing how the environment restricts both the production of flowers and the conversion of flowers into fruits.

The sandy valleys of central Amazonia have the highest abundance of monocot herbs (Costa 2006). Previous studies have suggested dispersal at short distances by insects (Horvitz 1981; Horvitz and Schemske 1994; Gómez and Espadaler 2013; Santana *et al*. 2016; Matlaga *et al*. 2017) and spatially dependent recruitment of seed sources (Rodrigues and Costa 2012) to be important causes for this limited distribution. Our results suggest the greater production of flowers and the greater availability of fruits (and seeds), in addition to the seed source-dependent recruitment at short distances, are the causal factors for the limited distribution of these plants in sandy soils.

### Synergistic effect of soil conditions and rainfall

Although flower and fruit production of monocot herbs were higher in sandy soils, rainfall also affects fruit production and how soil conditions affect fruit production (compare c and d in Fig. 3B). The higher abundance of herb species in sandier soils (Costa 2006; Drucker *et al*. 2008), which characterizes the bottom valleys in central Amazonia reflects their requirements (Tuomisto and Ruokolainen 1994; Tuomisto and Poulsen 1996). Requirements for reproduction and survival might include high humidity, poorly drained soils, suitable conditions for specific pollinators (Misiewicz *et al*. 2014) and dispersers (Bodmer 1991; Salas 1996; Fragoso 1997), more incidence of light (Castilho *et al*. 2006), and less herbivores (Pacheco 2001; Fine 2004; Fine *et al*. 2006).

Previous studies suggest the low fruit initiation might be correlated with insufficient pollen deposition (Ashman *et al*. 2004; Knight *et al*. 2005; Ågren *et al*. 2008) and pre-dispersal predation (Janzen 1971; Fernández *et al*. 2007; Boieiro *et al*. 2012; Jackson *et al*. 2022). In this sense, pollinators are less abundant in the rainiest months at the study site (Oliveira 1999) at the same time as herbivorous tend to increase in abundance and activity at the wetter months (Coley and Barone 1996; Baltzer and Davies 2012; Espelta *et al*. 2017).

The months with higher rainfall (> 250 mm) were not associated with greater flower production, and rainfall was negatively associated with the conversion of flowers to fruits at clayey sites. This is against expectations because hydric restrictions reduce the development of the male gametophyte (Saini 1997) and also increases the floral abortion rate (Akhalkatsi and Lösch 2005). However, the negative influence of rainfall on fruit production corroborates with studies that found no correlation between flower production and water supplementation (Ågren *et al*. 2008). These results may indicate that the amount of water in the soil is enough to supplement all water needed for reproduction, and rainfall might be a stressor factor rather than a positive push towards the establishment. Indeed, the mechanic impact of heavy rains and winds in wet season might increase the chance of physical damage by brach and tree fall (Brokaw 1985) and the probability of herbivorous attack (Baltzer and Davies 2012).

Rainfall in central Amazonia is relatively high even during the driest months compared to other forests (Ratter *et al*. 1997), and in these conditions, monocots herbs might not be directly affected by changes in rainfall to the same extent as to variation in edaphic conditions. In contrast, water availability can be constrained by distance to water sources (Chauvel *et al*. 1987; Rennó *et al*. 2008), and soil granulometry (Hodnett and Tomasella 2002), which might explain why the production of flowers and fruits was highest in sandy areas close to streams.

The change in the reproduction of herbaceous species along the sand gradients in central Amazonia might restrict species distribution and affect the composition of whole communities (Costa *et al*. 2005; Drucker *et al*. 2008; Rodrigues and Costa 2012; Schietti *et al*. 2014). During the rainy months, the difference in fruit production between the sandy lowlands and the clayey uplands become much more pronounced than in the dry months. Therefore, the more restricted distribution of monocot herbs to sandy areas might result from the higher production of fruits in these areas when rainfall is high. Rainfall is likely to reduce in central Amazonia under future climates scenarios (Costa and Pires 2010). Under this perspective and considering the reproductive restrictions along edaphic and climatic gradients presented here, the production of fruits in sandy areas might be reduced, and the abundance of herbaceous species in sandy valleys might become less pronounced.

## FUNDING

This work was supported by the Conselho Nacional de Desenvolvimento Científico e Tecnológico – CNPq [grant numbers 575637/2008-0, 473474/2008-5, 170183/2011-4].

ACKNOWLEDGMENTS

We thank the Programa de Pesquisas em Biodiversidade (PPBio) and INPA for the logistic support.

## SUPPLEMENTARY INFORMATION

The statistical model was compiled into R functions and a documented R script demonstrating its use is available in https://datadryad.org **[Supplementary Information S1]**.

## Notes

### Competing Interest Statement

The authors have declared no competing interest.

